# Non-uniform refinement: Adaptive regularization improves single particle cryo-EM reconstruction

**DOI:** 10.1101/2019.12.15.877092

**Authors:** Ali Punjani, Haowei Zhang, David J. Fleet

## Abstract

Single particle cryo-EM is a powerful method for studying proteins and other biological macromolecules. Many of these molecules comprise regions with varying structural properties including disorder, flexibility, and partial occupancy. These traits make computational 3D reconstruction from 2D images challenging. Detergent micelles and lipid nanodiscs, used to keep membrane proteins in solution, are common examples of locally disordered structures that can negatively affect existing iterative refinement algorithms which assume rigidity (or spatial uniformity). We introduce a cross-validation approach to derive *non-uniform refinement*, an algorithm that automatically regularizes 3D density maps during iterative refinement to account for spatial variability, yielding dramatically improved resolution and 3D map quality. We find that in common iterative refinement methods, regularization using spatially uniform filtering operations can simultaneously over- and under-regularize local regions of a 3D map. In contrast, *non-uniform refinement* removes noise in disordered regions while retaining signal useful for aligning particle images. Our results include state-of-the-art resolution 3D reconstructions of multiple membrane proteins with molecular weight as low as 90kDa. These results demonstrate that higher resolutions and improved 3D density map quality can be achieved even for small membrane proteins, an important use case for single particle cryo-EM, both in structural biology and drug discovery. *Non-uniform refinement* is implemented in the *cryoSPARC* software package and has already been used successfully in several notable structural studies.

## 1 Introduction

Single particle cryogenic electron microscopy (cryo-EM) is a rapidly advancing technique for determining high resolution 3D structures of proteins and protein complexes. There are several reasons that cryo-EM has become a tool of choice for recent structural biology projects [5], but key among them is the fact that cryo-EM can be used to study classes of proteins for which other techniques (e.g., X-ray crystallography and nuclear magnetic resonance) are not effective. These include membrane proteins, proteins larger than ∼30kDa with multiple conformational states, or with flexible or disordered domains. In all of these cases, the proteins are composed of regions with varying structural properties. Membrane proteins in particular are often encapsulated in detergent micelles or lipid nanodiscs, creating a large disordered region of density around the hydrophobic region of the protein. Despite the common occurrence of such spatial variability, most state-of-the-art 3D refinement algorithms are based on the mathematical assumption of uniformity (rigidity) of the particle.

This paper formulates a cross-validation regularization framework for single particle cryo-EM refinement, one that explicitly provides for the spatial inhomogeneity generated by various physical phenomena (disorder, motion, occupancy, etc) found in a typical molecular complex. Our proposed framework incorporates general domain knowledge about proteins, without specific knowledge of any particular molecule. Within this framework we derive a novel refinement algorithm, called *non-uniform refinement*, that accounts for structural variability, while taking care to ensure that the algorithm retains favourable statistical properties for validation and that it does not introduce avenues for over-fitting during 3D reconstruction.

With a GPU accelerated implementation of non-uniform refinement implemented in the *cryoSPARC* software package [22], we demonstrate improvements in resolution and 3D density map quality for a range of membrane proteins, yielding state-of-the-art reconstructions in all cases. We show results on a small membrane protein-Fab complex in a lipid nanodisc, a small membrane protein with negligible density outside of the lipid bilayer, a medium-sized membrane complex with a large detergent micelle, and a sodium channel with flexible domains. A beta version of the algorithm was previously released in *cryoSPARC* and has been used in structural studies of several other interesting proteins (see Section 8). Non-uniform refinement is reliable and automatic, requiring no change in parameters between datasets, and with no reliance on hand-made spatial masks or manual labels.

## 2 Regularization in Iterative Refinement

In the standard cryo-EM 3D reconstruction problem set up [12, 22, 26], a generative model describes the formation of images of a target protein in an electron microscope. According to the model, the unobserved protein structure, represented by a 3D density, is rotated by a 3D pose and projected along the direction of the electron beam. The resulting 2D projection is translated in the plane, modulated by the microscope CTF, and corrupted by additive noise, under a certain noise model. The 3D density map *m* is typically parameterized as a real-space 3D array with density at each voxel, in a Cartesian grid of box size *N*, and a corresponding discrete Fourier representation, 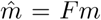. The goal of reconstruction is to infer the unobserved 3D densities of the voxel array, called **model parameters**. Representing the 3D density, 2D projections, observed images, and the noise model in the Fourier domain is common practice for computational efficiency, exploiting the well-known convolution and Fourier-slice theorems [1, 12, 22, 26]. The unobserved pose variables for each image are **latent variables**.

Iterative refinement methods (Algorithm 1), which provide state of the art results (e.g., [1, 11, 22, 35]), can be interpreted as variants of block-coordinate descent or the expectation-maximization algorithm [7], performing maximum-likelihood or maximum-*a-posteriori* estimation of model parameters from observations, according to the generative model described above. In cryo-EM, and more generally in inverse problems with noisy, partial observations, a critical component that modulates the quality of the results is regularization. Regularization, in intuitive terms, refers to the inclusion of prior knowledge to help penalize model complexity and avoid over-fitting. In cryo-EM refinement it plays a key role in mitigating the effects of noise, so that signal alone is used to infer model parameters. One can regularize problems explicitly, using a prior distribution over model parameters, or implicitly, by applying a regularization operator to the model parameters during optimization. Iterative refinement methods tend to use implicit regularizers, attenuating noise in the reconstructed map at each iteration. In either case, the separation of signal from noise is the crux of many inference problems.

### Algorithm 1 Iterative Refinement (Expectation-Maximization)

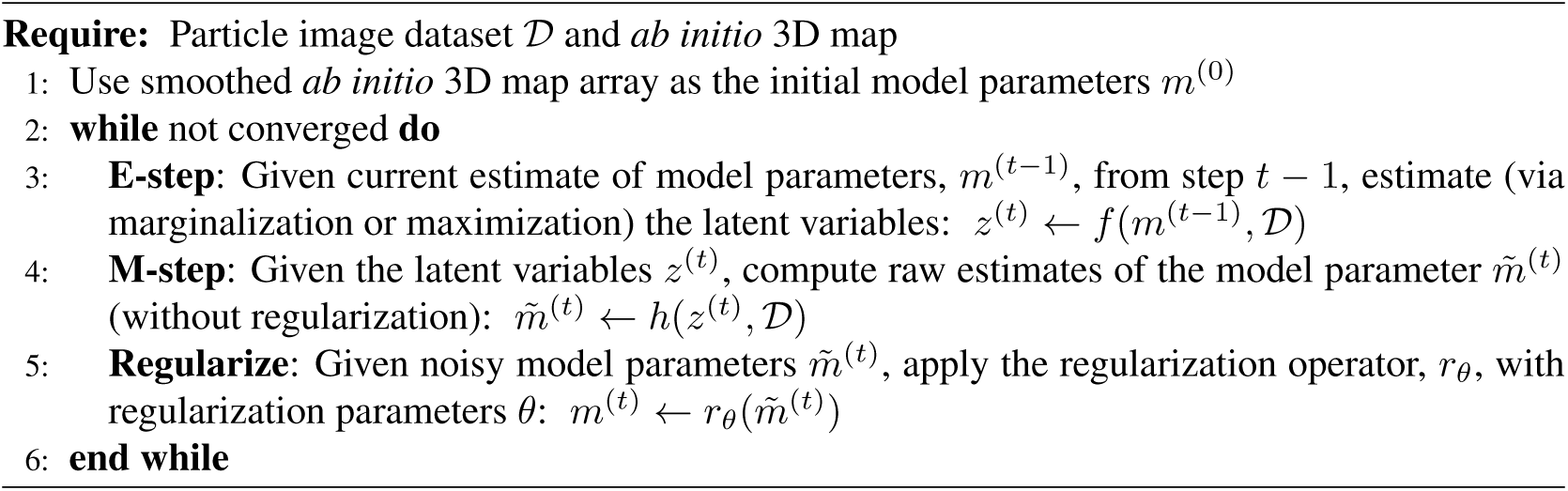

In the cryo-EM refinement problem, like many latent variable inverse problems, there is an additional interplay between regularization, noise buildup, and the estimation of latent variables. Retained noise due to under-regularization will contaminate the estimation of latent variables. This contamination is propagated to subsequent iterations and causes over-fitting. In Fig. 1, with the TRPA1 membrane protein [21], over-fitting occurs in the disordered micelle and in the dynamic tail at the bottom of the protein. Conversely, over-regularization attenuates useful signal structure, further degrading latent variable estimates. With TRPA1, signal loss occurs in the central region where the structure is not as well resolved as it otherwise could be. In general, the amount of regularization (or smoothing) depends on a set of regularization parameters, the tuning of which is often critical to successful outcomes.

**Figure 1:**
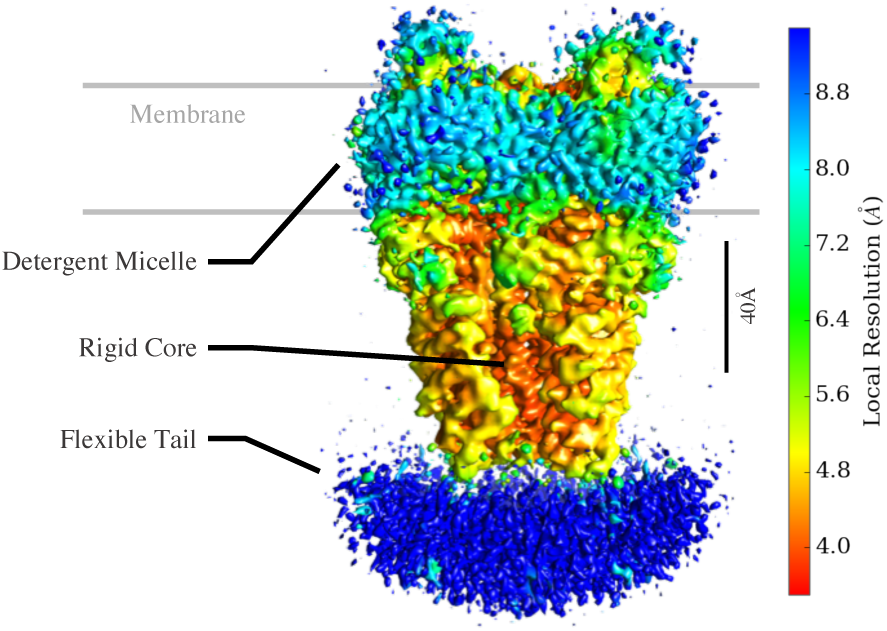
A 3D map from uniform iterative refinement (in *cryoSPARC*) reveals spatial variations in structure properties in a prototypical membrane protein (TRPA1 ion channel, EMPIAR-10024 [21]). The density map is shown with a low threshold, where color depicts local resolution [2] as a proxy for local structure properties. The core inner region (red) is more rigid and resolved to higher resolution. The solvent facing region (yellow) is less well ordered. The detergent micelle (light blue) is largely disordered. A flexible tail at the bottom (blue) is blurred due to motion. In uniform refinement, a shift-invariant regularizer smooths all regions equally. This uniformity leaves high frequency noise to accumulate in disordered regions (under-regularization), while discarding resolvable signal in rigid regions (over-regularization). Non-uniform refinement is designed to alleviate these problems.

This paper reconsiders the task of regularization based on the observation that common iterative refinement algorithms often systematically under-fit *and* over-fit different regions of a 3D structure, simultaneously. This causes a loss of resolvable detail in some parts of a structure, and accumulation of noise in others. The reason stems from the use of frequency-space filtering as a form of regularization. Some programs, like *cisTEM* [11], use a strict resolution cutoff, beyond which Fourier amplitudes are set to zero before alignment of particle images to the current 3D structure. In *RELION* [27], regularization was initially formulated with an explicit Gaussian prior on Fourier amplitudes of the 3D structure, with a hand-tuned parameter that controls Fourier amplitude shrinkage. Later versions of *RELION* [28] and *cryoSPARC*’s homogeneous (uniform) refinement [22] use a transfer function (or Wiener filter) determined by Fourier Shell Correlation (FSC) computed between two independent half-maps (e.g., [3, 13, 25]).

Such methods presume a Fourier basis and shift-invariance. Although well-suited to stationary processes, they are less well suited to protein structures, which are spatially compact and exhibit non-uniform disorder, motion, or variations in signal resolution. FSC, for instance, provides an aggregate measure of resolution. To the extent that FSC under- or over-estimates resolution in different regions, FSC-based shift-invariant filtering will over-smooth some regions, attenuating useful signal, and under smooth others, leaving unwanted noise. To address these issues we introduce a family of adaptive regularizers that can, in many cases, find better 3D structures with improved estimates of the latent poses during refinement.

## 3 Cross-Validation Regularization

We formulate a new regularizer for cryo-EM reconstruction in terms of the minimization of a cross-validation objective [10, 31]. Cross-validation (CV) is a general principle that is widely used in machine learning and statistics for model selection and parameter estimation with complex models. In CV, observed data are randomly partitioned into a training set and a held-out validation set. Model parameters are inferred using the training data, the quality of which is then assessed by measuring an error function applied to the validation data. In **k-fold CV**, the observations are partitioned into *k* parts. In each of *k* trials, one part is selected as the held-out validation set, and the remaining *k* − 1 parts comprise the training set. The per-trial validation errors are summed, providing the total CV error. This procedure measures agreement between the optimized model and the observations, without bias due to over-fitting. Rather, over-fitting during training is detected directly as an increase in the validation error. Importantly, formulating regularization in a cross-validation setting provides a principled way to design regularization operators that are more complex than the conventional, isotropic frequency-space filters. The CV framework is not restricted to a Fourier basis. One may consider more complex parameterizations, the use of meta-parameters, and incorporate cryo-EM domain knowledge.

Given a family of regularizers *r*_*θ*_ with parameter *θ*, the minimization of CV error to find *θ* is often applied as an *outer loop*. This requires the optimization of model parameters *m* to be repeated many times with different values of *θ*, a prohibitively expensive cost for problems like cryo-EM. Instead, one can also perform CV optimization as an *inner loop*, while optimization of model parameters occurs in the outer loop. Regularizer parameters *θ* are then effectively optimized on-the-fly, preventing under- or over-fitting without requiring multiple 3D refinements to be completed.

To that end, consider the use of 2-fold CV optimization to select the regularization operator, denoted *r*_*θ*_(*m*) in the regularization step in Algorithm 1 (note that *k* > 2 is also possible). The dataset 𝒟 is partitioned into two halves, 𝒟_1_ and 𝒟_2_, and two (unregularized) refinements are computed, namely *m*_1_ and *m*_2_. For each, one half of the data is the ‘training set’, and the other is held out for validation. To find the regularizer parameters *θ* we wish to minimize the total CV error *E*, i.e.,

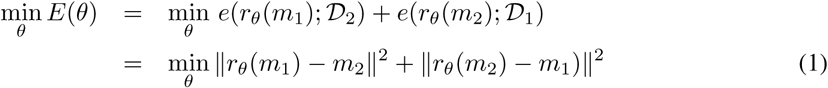

where *e* is the negative log likelihood of the validation half-set given the regularized half-map. The second line simplifies this expression by using the raw reconstruction from the opposite half-set as a proxy for the actual observed images. Note that assumptions for “gold standard” refinement [28] are not broken in this procedure (see Section 5). With the L2 norm, Eqn. 1 reduces to a sum of per-voxel squared errors, corresponding to white Gaussian noise between the half-set reconstructions. When the choice of *θ* causes *r*_*θ*_ to remove too little noise from the raw reconstruction, the residual error *E* will be unnecessarily large. If *θ* causes *r*_*θ*_ to over-regularize, removing too much structure from the raw reconstruction, then *E* increases as the structure retained by *r*_*θ*_ no longer cancels corresponding structure in the opposite half-map. As such, minimizing *E*(*θ*) provides the regularizer that optimally separates signal from noise. Similar objectives have been used for image de-noising [18]. Finally, we note that this formulation can be extended to compare each half-set reconstruction against images directly (dealing appropriately with the latent pose variables) or to use error functions corresponding to different noise models.

## 4 Regularization Parameter Optimization

The CV formulation in Eqn. 1 provides great flexibility in choosing the family of regularizers *r*_*θ*_, taking domain knowledge into account. For non-uniform refinement, we wish to accommodate protein structures with spatial variations in disorder, motion, and resolution. Accordingly, we define the regularizer to be a space-varying linear filter. The filter’s spatial extent is determined by the regularization parameter *θ*(*x*), which varies with spatial position:

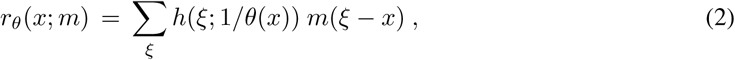

where *h*(*x*; *ψ*) is symmetric smoothing kernel, the spatial scale of which is specified by *ψ*. In practice we let *h*(*x*; *ψ*) be a 2^nd^-order Butterworth kernel, the parameter of which is the upper band-limit (frequency) of its transfer function. In this context *θ*(*x*) represents the corresponding wavelength.

When Eqn. 2 is combined with the CV objective for the estimation of *θ*(*x*), one obtains

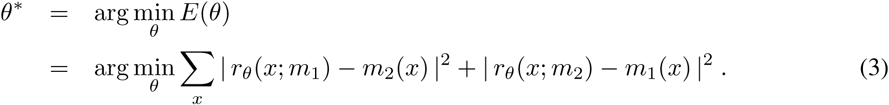

With one regularization parameter at each voxel, i.e., *θ*(*x*), this reduces to a large set of decoupled optimization problems, one for each voxel. That is, for voxel *x* one obtains

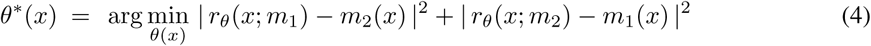

With this decoupling, *θ*(*x*) can transition quickly from voxel to voxel, yielding high spatial resolution. On the other hand, the individual sub-problems (4) are not well constrained since each parameter is estimated from data at a single location, so the parameter estimates are not useful. In essence, our regularizer design has two competing goals, namely, reliable signal detection, and high spatial resolution (i.e. respecting boundaries between regions with different properties). Signal detection improves through aggregation of observations (e.g., neighboring voxels), while high spatial resolution prefers minimal aggregation (as in Eqn. 4).

To improve signal detection, we further constrain *θ*\* to be smooth. That is, although in some regions *θ* should change quickly (solvent-protein boundaries), in most regions we expect it to change slowly (solvent and regions of rigid protein mass). Smoothness effectively limits the number of degrees of freedom in *θ*, which is important to ensure *θ* itself does not overfit during iterative refinement (see Sec. 7). One can encourage smoothness in *θ* by explicitly penalizing spatial derivatives of *θ* in the objective (Eqn. 3), but this yields a Markov random field problem that is hard to optimize. Alternatively, one can express *θ* in a low-dimensional basis (e.g., radial basis functions), but this requires prior knowledge of the expected degree of smoothness. Instead, we adopt a simple but effective approach. Assuming that *θ* is smoothly varying, we treat measurements in the local neighborhood of *x* as additional constraints on *θ*(*x*). A window function can be used to gives more weight to points close to *x*. We thereby obtain the following least-squares objective:

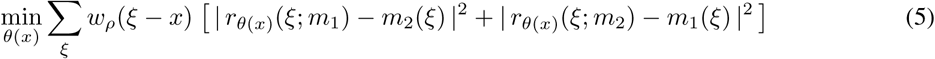

where *w*_*ρ*_(*x*) is positive and integrates to one, with spatial extent *ρ*. This allows one to estimate *θ* at each voxel independently, while the overlapping neighborhoods ensure that *θ*(*x*) varies smoothly.

### Algorithm 2 Regularization Step for Non-Uniform Refinement

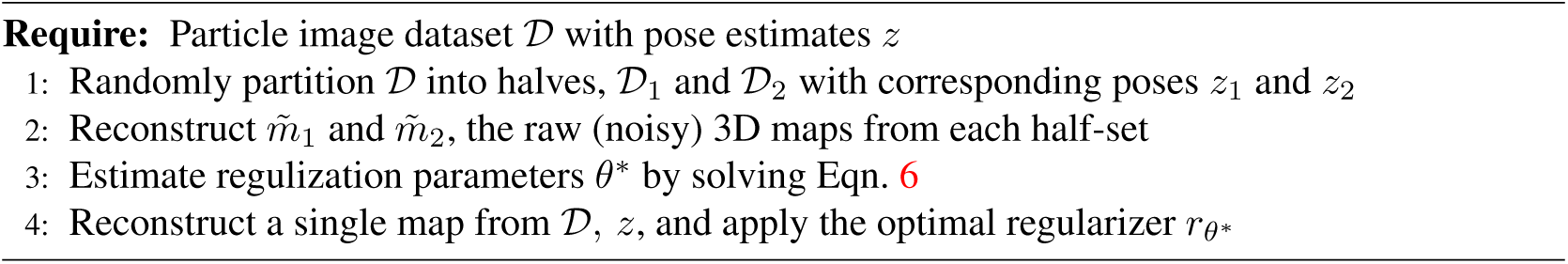

This approach also provides a natural way to allow for variable neighbourhood sizes, where *ρ*(*x*) depends on location *x*, so both rigid regions and transition regions are well modeled. Importantly, we want *ρ*(*x*) to be large enough to reliably estimate *θ*(*x*), but small enough to enable local transitions. A reasonable balance can be specified in terms of the highest frequency with significant power as captured by the regularization parameter *θ*(*x*). In particular, as a heuristic it suffices to ensure that *ρ*(*x*) > *γθ*(*x*) where *γ*, the adaptive window factor (AWF), is a constant.^1^ This constraint yields the final computational problem solved in non-uniform refinement to regularize 3D electron density at each iteration; i.e.,

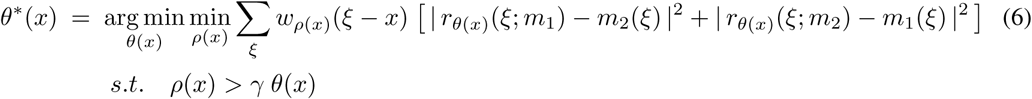

## 5 Non-Uniform Refinement Algorithm

Given a set of particle images and a low resolution *ab initio* 3D map, non-uniform refinement comprises three main steps, similar to conventional uniform (homogeneous) refinement (Alg. 1). The data are randomly partitioned into two halves, each of which is used to independently estimate a 3D half-map. This “gold standard” refinement [28] allows use of FSC for evaluating map quality, and for comparison with existing algorithms. The alignment of particle images against their respective half-maps, and the reconstruction of the raw 3D density map (the E and M steps in Alg. 1) are also identical to uniform refinement.

The difference between uniform and non-uniform refinement is in the regularization step. First, in non-uniform refinement, regularization is performed independently in the two half-maps. As such, the estimation of the spatial regularization parameters in Algorithm 2 effectively partitions each half-dataset into quarter-datasets, and we often refer to the raw reconstructions in Algorithm 2 as quarter-maps. The non-uniform refinements on half-maps are therefore entirely independent, satisfying the assumptions of a “gold-standard” refinement [28]. By contrast, conventional uniform refinement uses FSC between half-maps to determine regularization parameters at each iteration, thereby sharing masks and regularization parameters, both of which contaminate final FSC-based assessment because the two half-maps are no longer reconstructed independently.

Most importantly, non-uniform refinement uses Eqn. 6 to define the optimal parameters with which to regularize each half-set reconstruction at each refinement iteration. Figure 2 shows an example of the difference between uniform filtering (FSC-based) and the new CV-optimal regularizer used in non-uniform refinement. Uniform regularization removes signal and noise from all parts of the 3D map equally. Non-uniform regularization, on the other hand, removes more noise from disordered regions, while retaining the high-resolution signal in well-structured regions that is critical for aligning 2D particle images in the next iteration.

**Figure 2:**
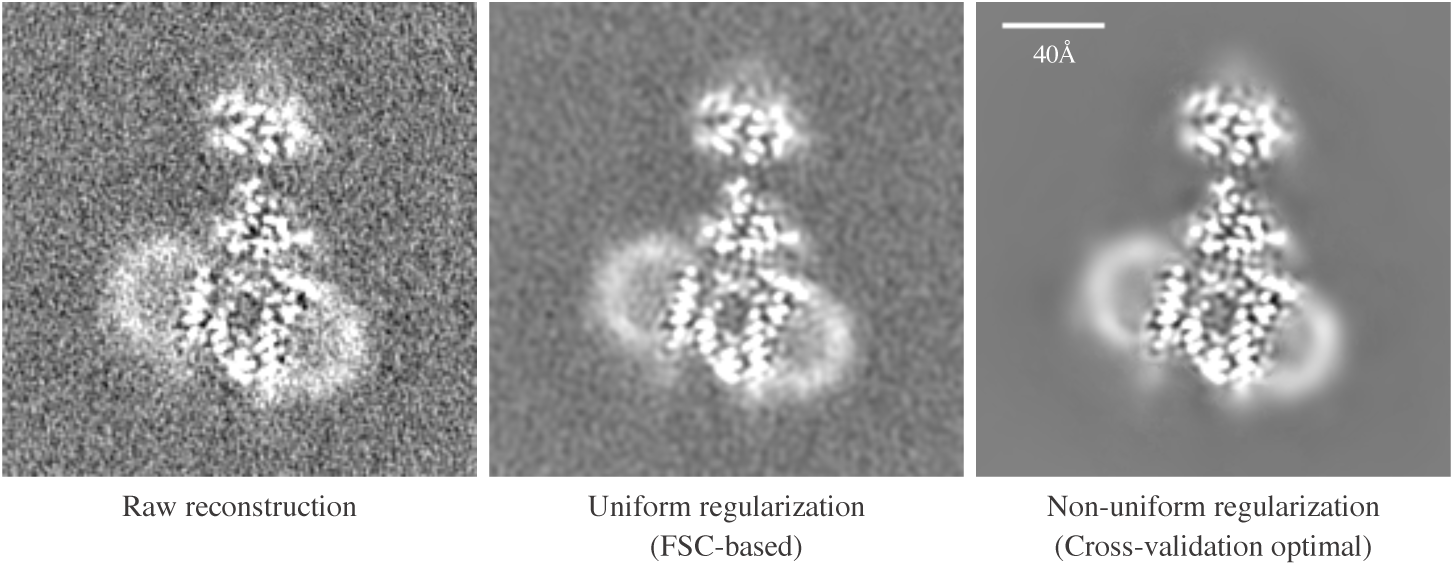
Images illustrate the difference between uniform (FSC-based) regularization and non-uniform (cross-validation-optimal) regularization on a membrane protein (PfCRT, EMPIAR-10330 [15]). **Left:** A central slice through a raw reconstruction (M-step of Alg. 1) of a half-map after 9 iterations. **Middle:** The same reconstruction is shown after uniform isotropic filtering (based on FSC between half-maps). **Right:** The same half-map reconstruction after non-uniform regularization with the optimal CV regularizer. Non-uniform regularization removes noise from the solvent background and nanodisc region, while preserving the high-resolution structure needed for particle alignments in the well-ordered protein region. It is particularly effective due to its implicitly small number of degrees of freedom (Eqn. 6) and capacity to model sharp transitions between ordered and disordered regions.

In practice, for regularization parameter estimation, Equation 6 is solved on a discretized parameter space where a relatively simple discrete search method can be used (e.g., as opposed to continuous gradient-based optimization). The algorithm is implemented in Python within the *cryoSPARC* software platform [22], with most of the computation implemented on GPU accelerators. An efficient solution to Eqn. 6 is important in practice because this subproblem is solved twice for each iteration of a non-uniform refinement.

Finally, the tuning parameters for non-uniform regularization are interpretable and relatively few in number. They include the order of the Butterworth kernel, the discretization of the parameter space, and the scalar relating *ρ*(*x*) and *θ*(*x*), called the adaptive window factor (AWF). In all experiments below we use a 2^nd^-order Butterworth filter, and a fixed AWF parameter *γ* = 2. We discretize the regularization parameters into 50 possible values, equispaced in the Fourier domain to provide greater sensitivity to small scale changes at finer resolutions. We find that non-uniform refinement is approximately two times slower than uniform refinement in our current implementation.

## 6 Results

Using four different membrane proteins as examples, we provide extensive comparisons between the proposed non-uniform refinement algorithm and conventional uniform (homogeneous) refinement. For the purposes of this comparison, all processing is performed in *cryoSPARC*. Except for the type of regularization used in refinement, all parameters and stages of data analysis are held constant between uniform and non-uniform refinement. Both begin with the same *ab-initio* structure. The same algorithm is used for estimating alignments between 2D particle images and 3D maps, and the same CTF-corrected back-projection is used for direct (unregularized) 3D reconstruction.

With each dataset the same mask is used for both uniform and non-uniform refinement when computing FSC curves. Masks are tested using phase randomization [3] to ensure they do not cause FSC bias. In addition, the same B-factor is used to sharpen both uniform and non-uniform refinement maps for each dataset. This consistency helps to ensure that the visible differences in 3D structure are due to algorithmic differences and not to differences in the amplification of high spatial frequencies. Where maps are colored by local resolution estimates, a straightforward implementation of *Blocres* [2] is used for resolution estimation.

Finally, all non-uniform refinement parameters (other than symmetry) are held constant across all datasets. No manual masks are used during alignment, no masking is done to identify or separate the micelle or nanodisc regions, and no modifications are made to the 2D particle images.

### 6.1 STRA6-CaM

Zebrafish STRA6-CaM [4] is a 180kDa C2 symmetric membrane protein complex comprising the 74kDa STRA6 protein bound to calmodulin (CaM). STRA6 mediates the uptake of retinol in various tissues. We present results of uniform and non-uniform refinement on a dataset of 28,848 particle images (pixel size

1.07*Å*) of STRA6-CaM in a lipid nanodisc, courtesy of Oliver Clarke and Filippo Mancia [6]. Note that this dataset differs from the older set used in [4] which consisted of 56,615 particle images of STRA6-CaM in amphipol.

Figure 3A shows FSC curves from uniform and non-uniform refinement, computed with the same mask. There is a substantial improvement in nominal resolution, from 4.0*Å* to 3.6*Å*. This indicates an improvement due to non-uniform refinement in the average global signal-to-noise over the entire structure. It is well known that different regions of a protein will have different resolution characteristics [2] and this is apparent in Figure 3C. The filtered and sharpened 3D maps from uniform and non-uniform refinements are shown from two viewing directions, colored by local resolution estimates. Both maps are filtered using their respective FSC curves and sharpened using a B-factor of − 140*Å*^2^. No local filtering or sharpening is used. The maps are thresholded to contain the same total volume in both cases. Non-uniform refinement resolves significantly improved structural detail in several regions of the molecule, while peripheral and flexible regions remain at low resolutions.

**Figure 3:**
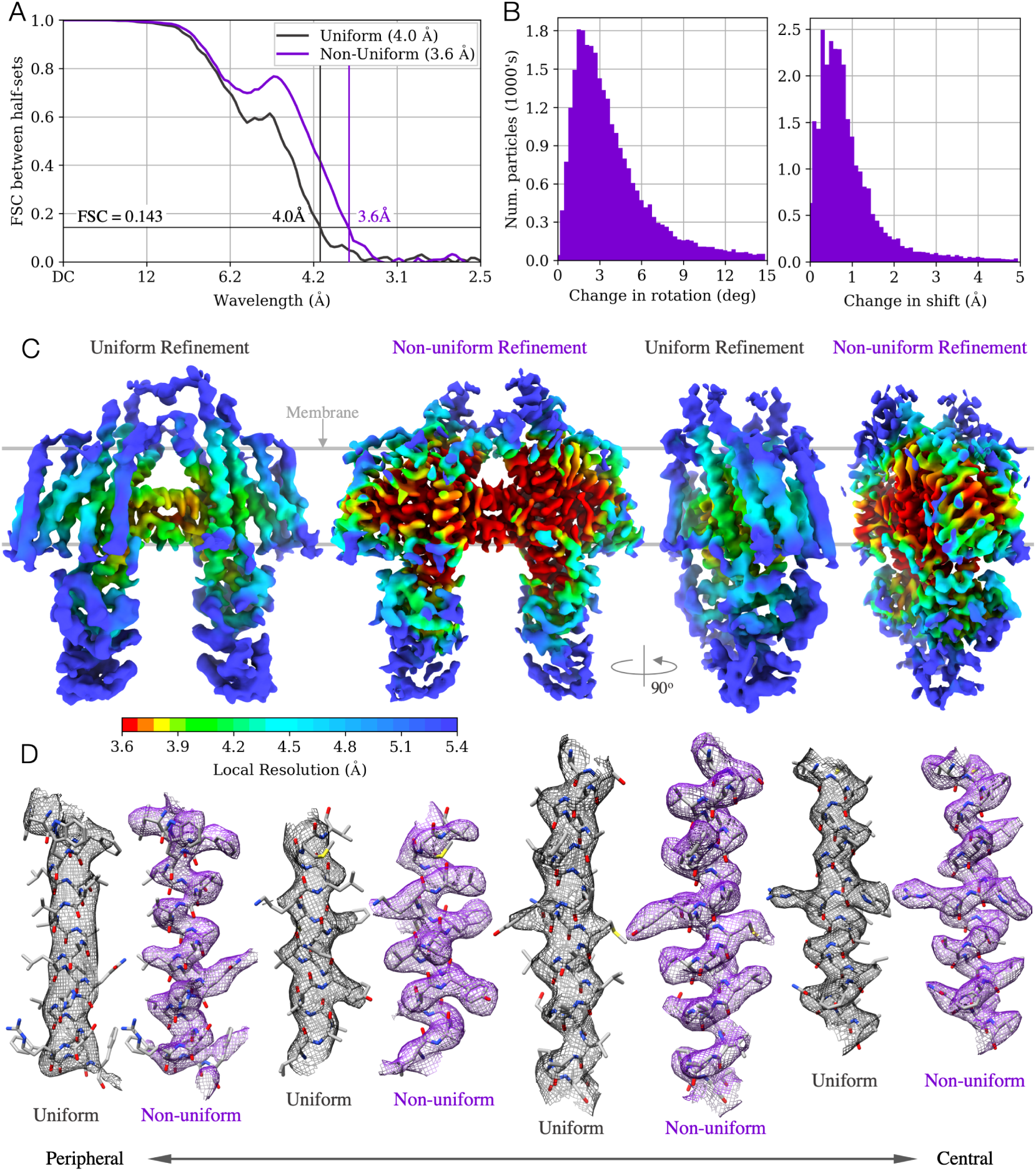
Results of uniform and non-uniform refinement from 28,848 particle images of **STRA6-CaM** [4] in lipid nanodisc. **A**: FSC curves computed with the same mask for both refinements. Numerical improvement from 4.0*Å* to 3.6*Å* reflects a spatially global improvement of signal. Improvement in some regions is even larger. **B**: Histograms of change in particle alignment pose and shift between uniform and non-uniform refinement. Optimal regularization in non-uniform refinement improves the ability to align particles over iterations. **C**: 3D density maps from uniform and non-uniform refinement are filtered using the corresponding FSC curves and sharpened with the same B-factor, − 140*Å*^2^. No local filtering or sharpening is used, and thresholds are set to keep the enclosed volume constant. Map differences are due to algorithmic rather than visualization differences. Map color depicts local resolution (*Blocres* [2]), on a single color scale. **D**: Individual *α*-helical segments from the non-uniform map (purple) resolve backbone and side-chains while the uniform map (grey) does not in most cases. The left-most *α*-helix is peripheral while the right-most is central.

Figure 3D shows detailed views of individual helices from within the structure with a docked atomic model, also courtesy of Oliver Clarke [6]. The visible difference in structure quality between uniform and non-uniform refinement is especially relevant during atomic model building, where in many cases, protein backbone and side-chains can only be traced with confidence in the non-uniform refinement map.

Non-uniform refinement appears to improve 3D reconstruction by improving particle image alignments. The STRA6-CaM particle is encased in a lipid nanodisc that has significant mass relative to the protein itself. Also, the CaM subunits are not completely rigid. The presence of this substantial disordered density breaks the assumptions behind uniform refinement. Noise is permitted to iteratively build up within the lipid nanodisc and flexible regions, while the core protein density is over-regularized and information useful for alignments is lost. In non-uniform refinement, the structure is optimally regularized according to cross-validation. The lipid nanodisc is smoothed while protein structure is maximally retained, leading to improved particle alignments. The change in particle image alignments between uniform and non-uniform refinement is shown in Figure 3B. The pose changes are particularly large, with most particles changing by more than 3 degrees. This change leads directly to improved structure quality.

In the original STRA6-CaM publication using amphipol, the dataset of 56,615 particles was refined to 3.9*Å* using a uniform refinement. In the result presented here, about half the number of particles (28,848) in lipid nanodisc yield a uniform refinement result of 4.0*Å* indicating that the quality of the data is higher. Non-uniform refinement, however, produces a 3.6*Å* result that is substantially better than both uniform refinements.

### 6.2 PfCRT

The *Plasmodium falciparum* chloroquine resistance transporter (PfCRT) is a small asymmetric 48kDa membrane protein with no soluble region [15]. Mutations in PfCRT are associated with the emergence of resistance to chloroquine and piperaquine as antimalarial treatments. The PfCRT dataset used here comprises 16,905 particle images of PfCRT in lipid nanodisc with a Fab bound, imaged with a 0.5175*Å* pixel size. It is available publicly as EMPIAR-10330 [15]. An early version of the non-uniform refinement algorithm was essential for reconstructing the published high resolution density map, into which a molecular model was built and from which biological insights were drawn [15].

For PfCRT, the difference in resolution and map quality between uniform and non-uniform refinement is dramatic. Figure 4A,C shows the corresponding FSC curves and 3D density maps. Uniform refinement is limited to 6.9*Å*, a resolution at which transmembrane helices are barely resolvable. At this low resolution, the authors [15] were unable to interpret the density map nor improve it with conventional techniques. By contrast, their results and ours here show that non-uniform refinement recovers signal up to 3.6*Å*, from which an atomic model can be built with confidence. Such levels of improvement are the difference between success and failure in a cryo-EM project.

**Figure 4:**
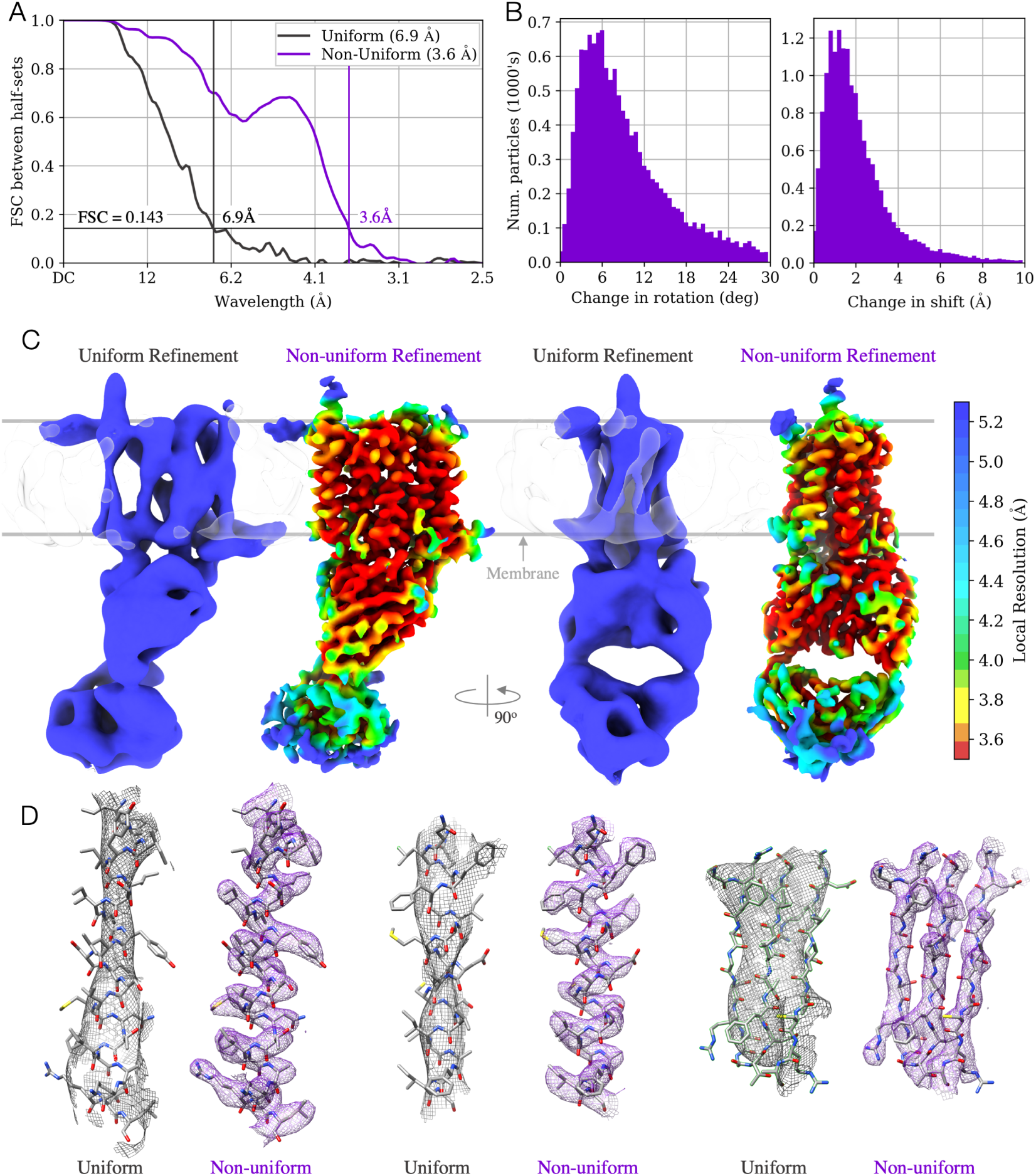
Results of uniform and non-uniform refinement from 16,905 particle images of **PfCRT** [15] in lipid nanodisc with a single Fab bound. **A**: FSC curves computed with the same mask show numerical improvement from 6.9*Å* to 3.6*Å*, a dramatic global improvement of signal. **B**: Histograms of change in particle alignments between uniform and non-uniform refinement. Optimal non-uniform regularization yields improved alignments through multiple iterations. Note that the *x*-axis limits are larger than in other figures. **C**: 3D density maps from uniform and non-uniform refinement, both filtered using the corresponding FSC curve, and sharpened with the same B-factor of − 100*Å*^2^. No local filtering or sharpening is used, and thresholds are set to keep the enclosed volume constant. Density differences are thus due to algorithmic rather than visualization differences. Maps are colored by local resolution from *Blocres* [2], all on the same color scale. **D**: Individual *α*-helical segments from the non-uniform map (purple) resolve backbone and side-chains while *α*-helices are barely resolved in uniform (grey) map density and *β*-strands are not separated.

The PfCRT-Fab complex (100kDa) is especially interesting due to its small size on the spectrum of proteins solvable by cryo-EM. In addition, the lipid nanodisc (∼50kDa) around PfCRT accounts for a large fraction of the total particle molecular weight. It is precisely in cases like this that the non-uniform refinement strategy is critical, as larger disordered regions leave more room for over- and under-regularization to influence particle alignments. For PfCRT in particluar, Fig. 4B shows substantial changes in particle alignments between uniform and non-uniform refinement. Most particle images exhibit pose changes of more than 6 degrees, indicating that regularization has a large impact on alignment. It also indicates that a large fraction of particle images may have been grossly misaligned by uniform refinement.

Figure 4D shows in detail the quality of the non-uniform refinement map for PfCRT. Transmembrane *α*-helices can be directly traced, including side-chains. In contrast, the uniform refinement map does not show helical pitch, and cannot separate *β*-strands in the Fab domain.

### 6.3 90kDa C3 Symmetric Membrane Protein

The next example is another small membrane protein with no soluble domain. This particle is C3 symmetric with a molecular weight of 90kDa, but unlike the PfCRT, does not have any Fabs bound. The protein mass is surrounded by a large detergent micelle, making alignments difficult. It is therefore a good test case for non-uniform alignment. The dataset, courtesy of Oliver Clarke [6], contains 42,740 particles at a pixel size of 1.05*Å*. It is part of an on-going study that is not yet published, so statistics and figures are shown here with permission, but we do not name the protein or show the entire 3D map.

Figure 5A shows FSC curves from uniform and non-uniform refinement. The overall global resolution of the map improves from 3.9*Å* to 3.6*Å* when using non-uniform refinement, but as with the STRA6-CaM dataset, local regions of the map improve in quality more than the numerical resolution improvement might suggest. Figure 5C shows densities for three transmembrane *α*-helices from the protein, with two (left, center) being closer to the periphery of the protein near the micelle, and one (right) being closer to the center. The peripheral *α*-helices show significant improvement in the level of detail resolved in the density map, allowing tracing of backbone and side-chains in the non-uniform refinement result. By comparison, model building would be difficult with the corresponding uniform refinement result. The improvement of the central *α*-helix is less dramatic, and was chosen to indicate that some parts of the uniform refinement map are resolved at its nominal 3.9*Å* resolution. Figure 5B highlights the large changes in particle alignment pose and shift between uniform and non-uniform refinement that also occur with this dataset.

**Figure 5:**
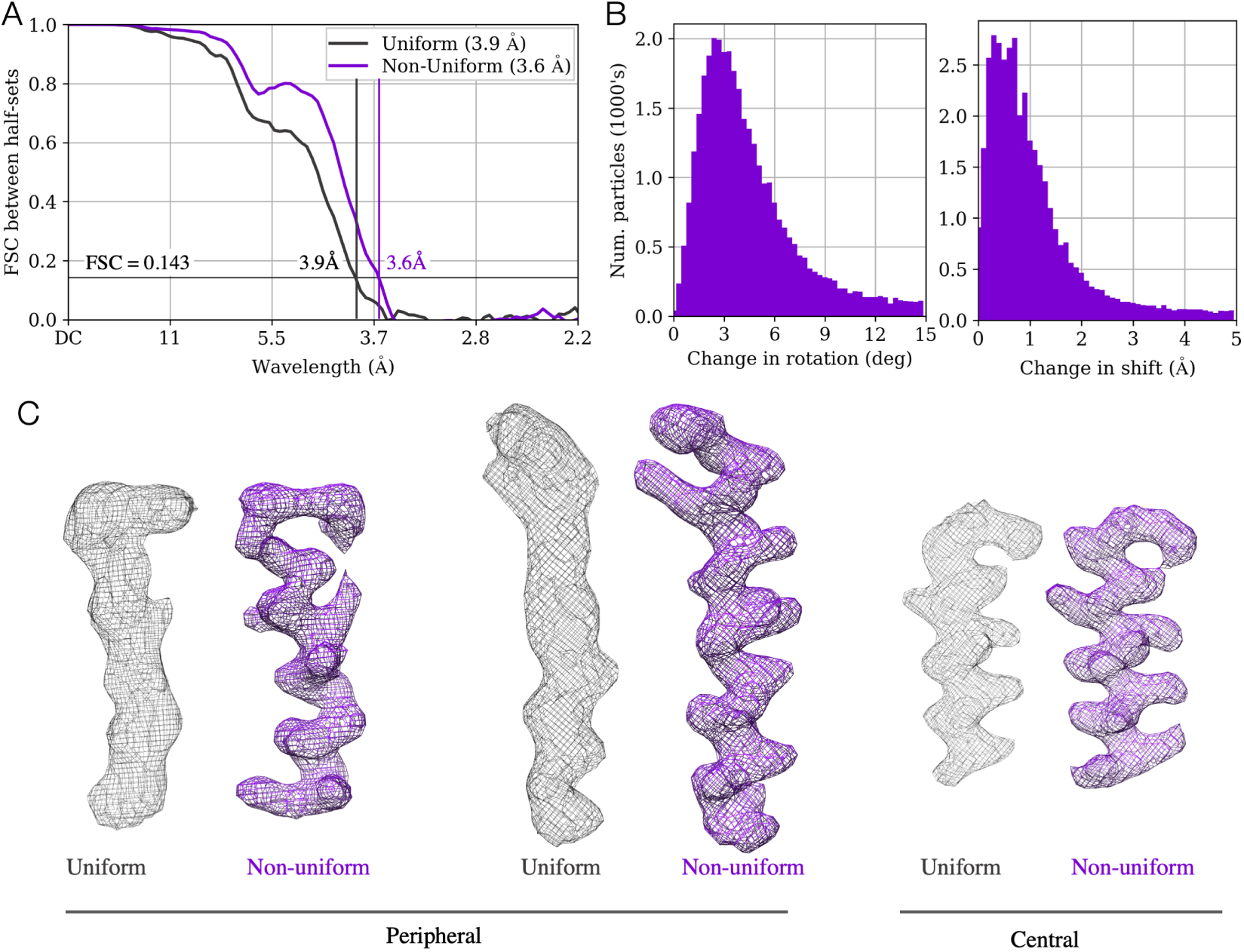
**Small membrane protein:** Results of uniform and non-uniform refinement on a dataset of 42,740 particle images of a C3 symmetric 90kDa membrane protein with no soluble domains [6]. **A**: FSC curves computed using the same mask show numerical improvement from 3.9*Å* to 3.6*Å*, indicating improved global average resolution. **B**: Histograms of change in particle alignments between uniform and non-uniform refinement. Optimal regularization in non-uniform refinement improves the ability to align particles over iterations, causing these changes. **D**: Density map detail for transmembrane *α*-helices. Two (left, center) are peripheral, near the micelle. One (right) is central within the protein. Both maps are filtered using the corresponding FSC curve, sharpened with the same B-factor of − 180*Å*^2^, no local filtering or sharpening is used, and thresholds are set to keep enclosed volume constant. Density differences are thus due to algorithmic rather than visualization differences. Non-uniform (purple) map has clear density for backbone and side-chains while for peripheral *α*-helices, helical pitch is barely resolved in uniform (grey) map density. The central *α*-helix (right) improves less dramatically.

### 6.4 Na_v_1.7 Ion Channel

The Na_v_1.7 channel [33] is a voltage-gated sodium channel found in the human nervous system. Nav channels are fundamental to the generation and conduction of action potentials in neurons, are mutated in various diseases, and are targeted by toxins and therapeutic drugs (e.g., for pain relief). We present results of uniform and non-uniform refinement on Na_v_1.7 data originally collected and processed by the authors of [33], available publicly as raw microscope movies (EMPIAR-10261). In this dataset, the Na_v_1.7 channel is bound to two Fabs, resulting in a 245kDa C2 symmetric complex solublized in detergent. The data contains 25,084 movies from a Gatan K2 Summit direct electron detector camera in counting mode, with a pixel size of 0.849*Å*. We processed the entire dataset through motion correction, CTF estimation, particle picking, 2D classification, *ab-initio* reconstruction and heterogeneous refinement in *cryoSPARC* v2.11 [22]. A total of 738,436 particle images were extracted from the movies. These were curated using 2D classification, yielding 431,741 particle images. As in [33], we detected two discrete conformational states corresponding to the active and inactive states of the channel. 130,982 particles were found in the inactive state, and the remaining 300,759 were found to be in the active state. We obtained state-of-the-art resolution reconstructions for both the active and inactive states, but here we focus solely on the active state.

Compared to the preceding datasets, the Na_v_1.7 complex has a larger molecular weight, and in relative terms, a smaller detergent micelle. It does, however, have other regions that are disordered or flexible, namely, a central 4-helix bundle, peripheral transmembrane domains, and the Fabs. Notably, non-uniform refinement handles these regions of the structure automatically along with the micelle, and therefore still delivers improved resolution and map quality. Figure 6A shows FSC curves from uniform and non-uniform refinement. Uniform refinement already reaches a 3.4*Å* resolution, and non-uniform refinement further improves this to 3.1*Å*. Figure 6C shows 3D density maps colored by local resolution. With non-uniform refinement, map quality is clearly improved in central transmembrane regions, while flexible parts of the structure (membrane *α*-helices closest to the micelle, Fab domains, and 4-helix bundle) remain at intermediate resolutions.

**Figure 6:**
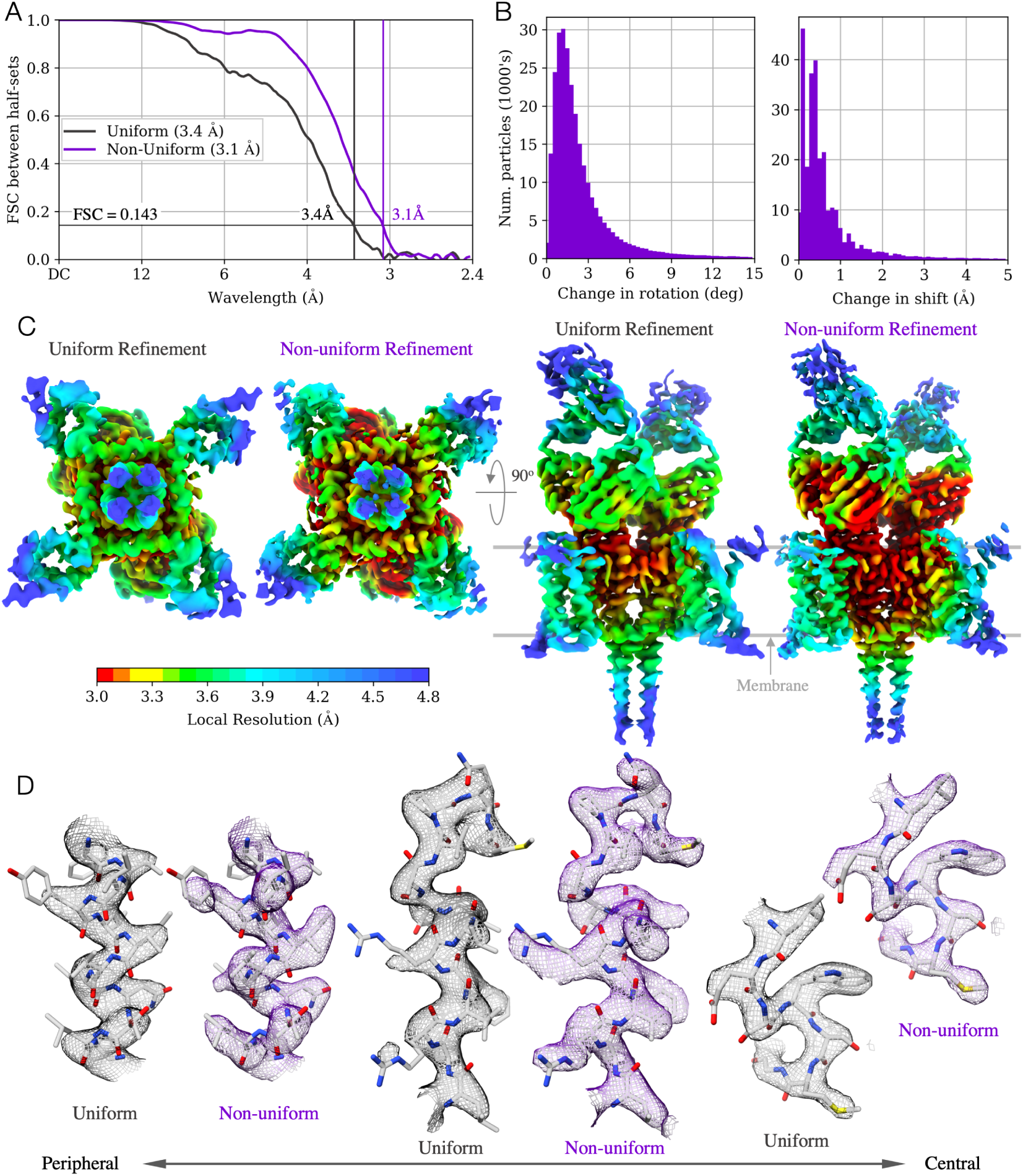
Results of uniform and non-uniform refinement on 300,759 particle images of **Na**_**v**_**1.7** [33] in a detergent micelle with two Fabs bound. **A**: FSC curves indicate significant improvement in signal with non-uniform refinement, with numerical improvement from 3.4*Å* to 3.1*Å*. **B**: Histograms show the changes in particle alignments enabled by optimal regularization in non-uniform refinement. Changes are somewhat smaller here compared to earlier examples. **C**: The 3D density maps are filtered using corresponding FSC curves, sharpened with a B-factor of − 85*Å*^2^. Thresholds are set to keep the enclosed volume constant. No local filtering or sharpening is used. Color depicts local resolution (*Blocres* [2]), on the same color scale. **D**: Individual *α*-helical segments from the non-uniform map (purple) are better resolved, allowing modelling of side-chain rotamers with confidence while uniform map (grey) often does not. Two *α*-helices (left, center), from the peripheral region of the transmembrane domain, show larger improvement. The other (right), from the central core, is already well resolved by uniform refinement.

Figure 6B shows changes in particle alignment poses and shifts between the two refinement types. These changes are smaller on average than the other examples, congruent with the smaller changes in map features. Figure 6D shows three selected regions from the density maps along with a docked atomic model (PDB-6N4Q [33]). In *α*-helices farther from the center of the protein (left, center), improvement in map quality is readily apparent, with many side chains being somewhat buildable in the uniform refinement map, but having clearly interpretable density in the non-uniform refinement map. Central *α*-helices (right) show less improvement, but map quality remains equal or slightly improved. This result is significant because it shows that reconstructions of proteins without disorder are not harmed by using non-uniform refinement.

Note that our result on the active state of the Na_v_1.7 channel is also an improvement over the published result of 3.6*Å* (EMDB-0341) [33]. In [33], the authors performed all processing in *cisTEM* [11]. In our result, a substantial part of the improvement over the published result is due to non-uniform refinement, while some other part may be due to improvements in other components of the processing pipeline in *cryoSPARC*.

### 6.5 Reconstruction B-factor Improvements

The preceding results indicate that non-uniform refinement improves particle image alignments during iterative refinement, yielding improved map resolution and quality. Here, we show that it also improves B-factors [25] of 3D reconstructions, indicating a more efficient use of signal from the same particle image data.

In [25], a B-factor is fit to the rotationally-averaged power spectrum of the reconstructed density map. The B-factor is composed of two parts, *B*_image_ and *B*_computation_. *B*_image_ captures the decay of high-frequency signal intrinsic to the imaging process. *B*_computation_ captures the additional high-frequency signal lost due to computational errors in alignment or reconstruction from the image data.

Figure 7A (left) shows overall B-factors fit to reconstructions for the four datasets discussed above, with uniform and non-uniform refinement, given the same image data. The Guinier plot in Fig. 7 (right) illustrates the difference in high-frequency signal decay between the two types of refinement, here for the Na_v_1.7 channel dataset. For all datasets, non-uniform refinement B-factor magnitudes are smaller than for uniform refinement. Since *B*_image_ is the same in both methods, it follows that non-uniform refinement has lower *B*_computation_, losing less high frequency signal to misalignment. This difference also suggests that using non-uniform refinement should in general require less image data to reach the same resolutions as uniform refinement, consistent with what we observed with the STRA6-CaM dataset above.

**Figure 7:**
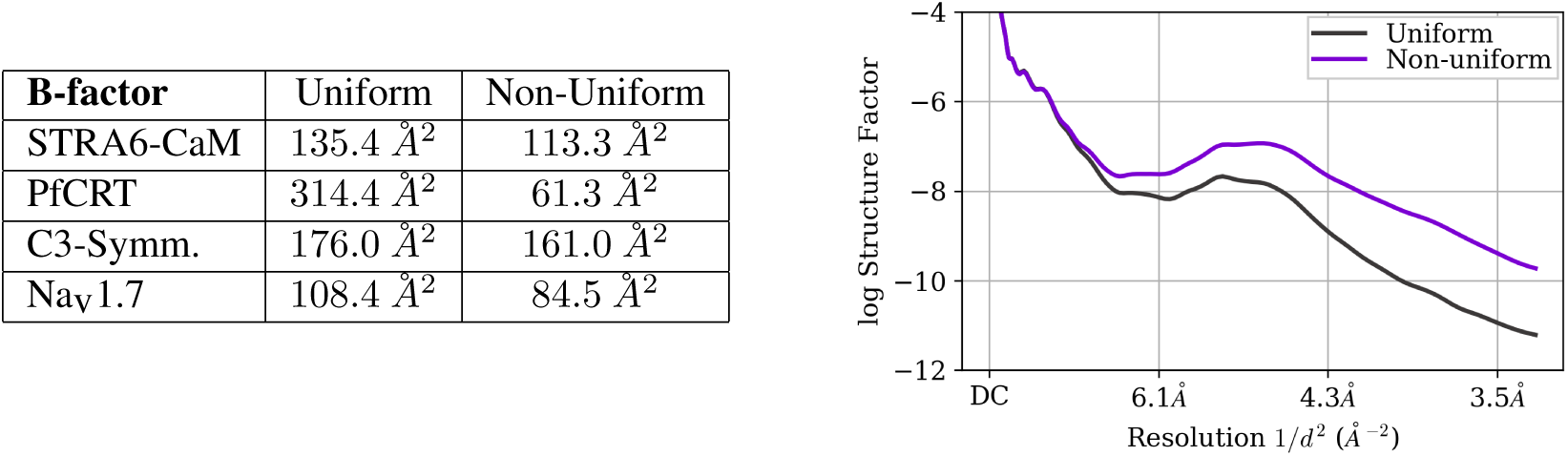
**Left**: B-factor values for four proteins for uniform and non-uniform refinement computed by *cryoSPARC*. Larger magnitudes (steep slopes) indicate greater signal lost. Given identical image data, differences in B-factor between uniform and non-uniform refinement indicate relative amounts of signal lost due solely to refinement. Non-uniform refinement generally has a significantly lower magnitude, indicating less less signal lost to noise in reconstruction at higher frequencies. **Right**: Guinier plot (log amplitude per frequency squared [25]) for the Na_v_1.7 channel dataset described in Sec. 6.4.

## 7 Discussion of Related Methods

There is a long line of cryo-EM research aimed at estimating the local resolution at each voxel of a 3D structure once a refinement is complete [2, 17, 29]. These methods are generally based on statistical tests applied to the coefficients of a windowed Fourier transform [2] or some form of wavelet transform [17, 29]. Although the aim of non-uniform refinement is not to estimate resolution *per se*, the regularizer parameter *θ*(*x*) does correspond to a local frequency band-limit at each voxel. As such, it might be viewed as a proxy for local resolution, but with some important differences. Notably, our formulation defines *θ*(*x*) as the optimum of a cross-validation objective that removes noise. It does not depend on a particular definition of “local resolution” nor on an explicit resolution estimator.

Local resolution estimates [8, 2] or local statistical tests [23] can also be used to adaptively filter a 3D map. This technique is used extensively for visualization, for assessing map quality in different regions of a particle, and during molecular model building. The family of filters and local filter band-limits are typically selected to maximize subjective visual quality and are therefore not optimized against the estimator used to determine local resolution. Considerations are also not made in this technique to control the number of degrees of freedom or model complexity of the local filter parameters. Furthermore, information from opposite half-set reconstructions is typically shared, breaking independence. While local resolution estimation followed by local filtering is generally satisfactory as a one-time post-processing step for visualization, in our experience it can lead to severe over-fitting when used iteratively in a refinement as a substitute for the regularization step in Alg. 1. Figure 8 gives an example of the over-fitting that often occurs. During iterative refinement, small mis-estimations of local resolution at a few locations (due to high estimator variance [2]) cause subtle over- or under-fitting, leaving slight density variations. Over multiple iterations of refinement, these errors can produce strong erroneous density that contaminate particle alignments and the local estimation of resolution itself, creating a vicious cycle. A related technique using iterative local resolution and filtering was described briefly in *EMAN2.2* documentation [1] and may suffer the same problem. The resulting artefacts (e.g., streaking and spikey density radiating from the structure) are particularly prevalent in datasets with junk particles, structured outliers, or small particles that are already difficult to align. To mitigate these problems, the approach we advocate couples an implicit resolution measure to a particular choice of local regularizer, with optimization explicitly designed to control model capacity and avoid over-fitting of regularizer parameters.

**Figure 8:**
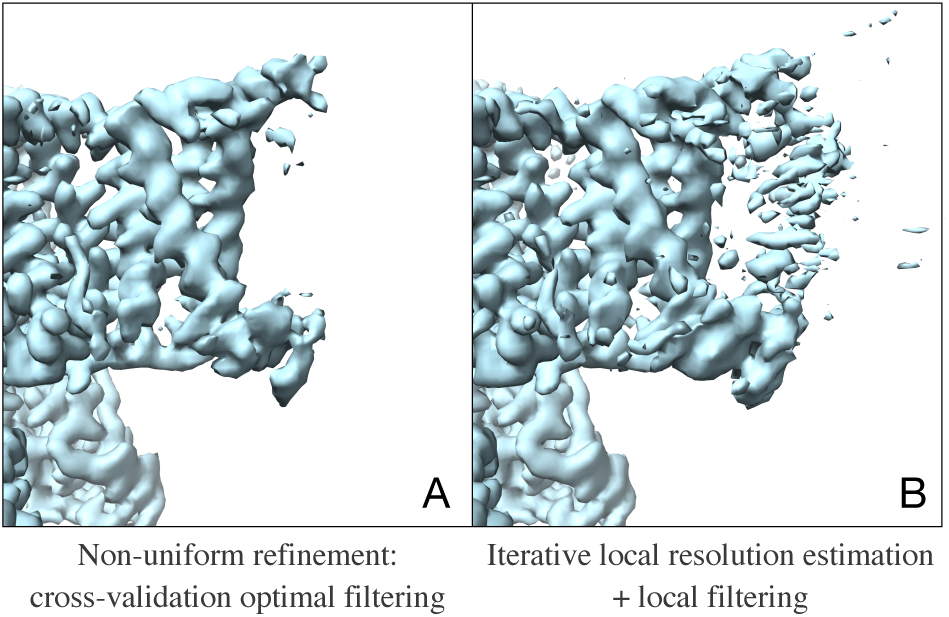
Side view of a peripheral transmembrane region of the Na_v_1.7 channel showing that naive forms of local filtering during iterative refinement can lead to significant over-fitting. **A**: Unsharpened density map at the 8^th^ iteration of non-uniform refinement as presented in this work. **B**: Unsharpened density map at the 8^th^ iteration of a simpler iterative refinement scheme where local resolution estimation and local filtering are performed at each iteration. Both maps are thresholded at the same level. The map in **B** has acquired clear artefacts of over-fitting, with incorrect high-resolution density dominating the peripheral transmembrane region of the protein. This artefactual density continues to strengthen over subsequent iterations, contaminating local resolution estimates and leading to errors in particle alignments.

Another related technique, used in *cisTEM* [11] and its predecessor *Frealign* [12], entails the manual creation of a mask to directly label a region of a structure that one expects to be disordered (eg., a detergent micelle), followed by low-pass filtering in that region to a single pre-set resolution at each iteration of refinement. This technique, one of the first to acknowledge and address the issue of simultaneous under- and over-fitting, shares the same intuitions behind non-uniform refinement. Nevertheless, it relies on manual decisions about how to regularize a map during refinement. Refinement is sensitive to regularization, and manual choices necessitate a trial and error process that can be tedious and difficult to replicate across datasets or by others.

Another possible approach to local control of regularization is to formulate the problem in another basis. In most single particle EM reconstruction methods, the Fourier basis is chosen primarily because the Fourier Slice Theorem facilitates fast projection evaluation and reconstruction. Another natural choice are the closely related spherical harmonics that permit fast rotation [34]. Wavelet bases have also been suggested [30], the basis functions of which are simultaneously localized in space and the Fourier domain. Indeed, wavelets are used in methods for local resolution estimation [2, 17, 29]. Kucukelbir et al [16] describe an interesting approach to iterative refinement with an adaptive wavelet basis and a sparsity prior. This approch is also similar in spirit to the goal in this work, but their model has a single regularization parameter for the entire 3D map. It is based on the noise variance estimated from the corners of particle images, which may not capture local variations in noise due to disorder, motion, or partial occupancy. Nevertheless, like the cross-validation approach here, the authors also find that with a data-driven approach they can largely avoid the need for manual parameter setting and the spatial masking that is needed with a Fourier basis.

## 8 Conclusions

We introduce non-uniform refinement, a new algorithm for achieving higher resolution and higher quality reconstructions of protein structures from single particle cryo-EM data. Non-uniform refinement uses cross-validation regularization as a principled tool for parameter estimation in noisy inverse problems like cryo-EM reconstruction. In this way, we are able to incorporate into the algorithm, and exploit, the domain knowledge that many types of proteins contain regions with varying levels of disorder, flexibility, or occupancy. These variations lead uniform refinement algorithms to under- or over-regularize regions of a 3D density map during refinement. Instead, non-uniform refinement automatically identifies and treats local regions optimally, without introducing bias into “gold standard” estimation of resolution with the FSC and without any specific knowledge about any particular protein.

A range of different proteins are used to show the capability of the algorithm to resolve higher-resolution detail, and in many cases, significantly improve map quality. A further effect of using a principled cross-validation procedure is that the entire algorithm becomes automatic and does not require user interaction to change any parameters between datasets.

In this work, non-uniform refinement is contrasted with uniform refinement in which global FSC-based resolution estimates are used for filtering during iterative refinement. The fact that non-uniform refinement locally regularizes 3D maps during refinement indicates that there may in fact be significant structure resolved beyond the global FSC-based resolution to which final maps are filtered before visualization. We do not address this possibility here, but it does raise the interesting question of how one can then effectively measure resolution in general (similar to the premise of [2]).

To construct the non-uniform refinement algorithm in this work, we have considered only one specific family of regularizers. It would be possible, within the same framework, to consider other families that may regularize even more effectively. For example, one could look to the extensive denoising literature for different regularizers.

In this paper, we have focused on results demonstrating the applicability of non-uniform refinement to membrane proteins, which are a large class with especially high importance in drug discovery. The algorithm, however, is general, and provides a useful approach for other proteins with local variation in physical properties.

A beta version of the non-uniform refinement algorithm was released in previous versions of *cryoSPARC* [22]. Indeed, it has been already been successful in helping to resolve a number of interesting new structures. Examples include a structure of FACT manipulating the nucleasome [19], multiple conformations of an ABC exporter [14], mTORC1 docked on the lysosome [24], the respiratory syncytial virus polymerase complex [9], a GPCR-G protein-*β*-arrestin megacomplex [20], and MERS-CoV/SARS-CoV with neutralizing antibodies [32]. An updated non-uniform refinement algorithm implementation, as described in this work, will be released in an upcoming version of *cryoSPARC* (www.cryosparc.com).

## Acknowledgements

We are extraordinarily grateful to Oliver Clarke and Yong Zi Tan for providing valuable cryo-EM data early in this project, and for sharing their experience as early adopters of non-uniform refinement and cryoSPARC. We thank the entire team at Structura Biotechnology Inc. that designs, develops, and maintains the cryoSPARC software system in which this project was implemented and tested. We thank John Rubinstein for comments on the manuscript. This work was financially supported in part by NSERC Canada and by the Canadian Institute for Advanced Research.

## Footnotes

1 Two key factors directly influence the choice of *ρ*(*x*) for estimating *θ*(*x*) reliably. The first is noise in the raw reconstruction. Aggregating squared error measures the residual variance (the residual power); larger windows provide more accurate variance estimates. The second is residual signal structure. When over-regularized, some high-frequency signal is lost, in which case signal residual is expected to have significant amplitudes near the wavelength *θ*. (Higher frequencies with insignificant amplitudes are dominated by noise, while lower frequencies are left unchanged by the regularizer and thus cancel the signal in the opposite half-map.) Integrating a squared band-pass signal provides a phase-independent measure of the local signal power; in our case the minimum window *ρ*(*x*) is determined by pass-band of the residual signal, which is expected to be close to *θ*. Thus the minimum *ρ*(*x*) is naturally constrained to be small multiple of *θ*.

